# LUNAR: A Deep Learning Model to Predict Glioma Recurrence Using Integrated Genomic and Clinical Data

**DOI:** 10.1101/2024.10.23.619762

**Authors:** Jessica A. Patricoski-Chavez, Seema Nagpal, Ritambhara Singh, Jeremy L. Warner, Ece D. Gamsiz Uzun

## Abstract

Gliomas account for approximately 25.5% of all primary brain and central nervous system tumors, with a striking 80.8% of these being malignant. The prognosis varies significantly; low-grade gliomas (LGGs) can exhibit 5-year survival rates of up to 80%, while higher-grade gliomas (HGGs) often see rates below 5%. Recurrence is a common challenge, occurring in 52%-62% of LGGs and 90% of HGGs, complicating clinical management and treatment planning. Currently, no widely available models exist for predicting glioma recurrence, which is critical for optimizing patient outcomes. Machine learning (ML) and deep learning (DL) techniques have shown promise in predicting recurrence for various cancers, most using Electronic Health Records (EHR). This study introduces g**L**ioma rec**U**rre**N**ce **A**ttention-based classifie**R** (LUNAR), a DL-based model to predict early versus late glioma recurrence by integrating clinical, genomic, and mRNA expression data from patients with primary grade II-IV gliomas. The data was obtained from The Cancer Genome Atlas and the Glioma Longitudinal Analysis Consortium (GLASS).

## 1. Introduction

Gliomas represent approximately 25.5% of all primary brain and central nervous system (CNS) tumors and 80.8% of malignant brain and CNS tumors.^1^ Gliomas are highly infiltrative tumors classified and graded based on molecular and genetic markers, degree of proliferation, and necrosis.^2^ Regardless of grade, gliomas are frequently characterized by their developed resistance to surgical and chemoradiation treatment regimens.^3,4^ The heterogeneity of glioma leads to wide variations in outcomes and prognosis, with low-grade gliomas (LGGs) reaching 5-year survival rates up to 80% while higher-grade glioma’s (HGGs) 5-year survival rate falls below 5%.^1^ The invasive nature of these tumors lends itself to a high likelihood of cancer recurrence, with 52%-62% of LGGs^5–7^ and 90% of HGGs^8,9^ recurring. Glioma recurrence occurs as early as 6 months to as late as 15 years.^5^ For all glioma types, recurrence poses a significant challenge to clinical management and treatment planning. Insight into a patient’s likelihood of recurrence can profoundly impact patient outcomes by optimizing intervention selection and timing, limiting unnecessary testing and procedures, and minimizing treatment-related disability.^10–14^ However, there are currently no widely available prediction models for assessing the risk of early glioma recurrence.

The numerous benefits of predicting cancer recurrence are not exclusive to glioma. As such, machine learning (ML) and deep learning (DL) methods have been applied to recurrence prediction tasks in multiple cancer types. González-Castro et al.^15^ evaluated the capacity of five ML classifiers to predict 5-year breast cancer recurrence using electronic health records. Their extreme gradient boosting model achieved an area under the receiver operating characteristic curve (AUROC) of 0.807, outperforming logistic regression, decision tree, gradient boosting, and multi-layer perceptron. Zuo et al.^16^ evaluated eleven different ML models for breast cancer prediction using clinical features and laboratory indices. Their highest performing model, Adaboost, achieved an AUROC of 0.987. Kumar et al.^17^ utilized dual convolutional neural networks (CNNs) to predict prostate cancer recurrence after radical prostatectomy using tissue images and achieved an AUROC of 0.81. Piedimonte et al.^18^ developed two ML models and two NNs to predict recurrence and time to recurrence in high-grade endometrial cancer and achieved a maximum AUROC of 71.8% with random forest using clinical data. In the case of glioma, Luo et al.^19^ successfully applied deep learning to predict glioma recurrence at multiple time points using clinical data and hematoxylin-eosin (H&E) stained slide images. Gonzalez-Bosquet et al.^20^ integrated clinical features and RNA-seq data from TCGA to predict endometrioid-type endometrial cancer recurrence. After testing over 170 ML models, they found their best model used only long non-coding RNA data and achieved an AUROC of 0.9.

Attention mechanisms have become increasingly prevalent across a wide range of DL applications^21^ Lan et al.^22^ developed DeepKEGG, which uses a biological hierarchy and self-attention to predict the recurrence of breast, liver, bladder, and prostate cancer, and reported AUROCs ranging from 0.799 to 0.961. Ai et al.^23^ developed a recurrence prediction model for non-small cell lung cancer using self-attention and CT images. Wang et al.^24^ developed hepatocellular carcinoma early recurrence prediction models that utilized intra- and inter-phase attention on clinical data, CT images, or both. Their model achieved a prediction accuracy of 81.2% and AUROC of 0.869. While the existing models are valuable, there is a current lack of glioma recurrence prediction models that incorporate both clinical and genomic data. Additionally, current models tend to rely on large datasets or imaging features, both of which are often difficult to procure.

Given this unmet need and recent successful applications of attention mechanisms to disease-classification tasks, we developed LUNAR, a novel g**L**ioma rec**U**rre**N**ce **A**ttention-based classifie**R**, to predict early vs. late glioma recurrence using clinical, mutation, and mRNA-expression data from patients with primary grade II-IV gliomas from The Cancer Genome Atlas^25^ and the Glioma Longitudinal Analysis Consortium.^26^

## 2. Materials and Methods

### 2.1. Datasets

The Cancer Genome Atlas (TCGA) has molecularly characterized tumors from over 11,000 patients across 33 cancer types, including multiple types of glioma.^25^ To create a robust dataset of all TCGA primary diffuse gliomas, we downloaded clinical, mutation, and gene expression data for the LGG and Glioblastoma Multiforme (GBM) merged dataset, GBMLGG,^27^ from cBioPortal (cbioportal.org)^28^ and University of California Santa Cruz (UCSC) Xena (xenabrowser.net).^29^ To minimize clinical data missingness, we supplemented the GBMLGG clinical data with clinical data from the TCGA LGG^30^ and TCGA GBM^31,32^ datasets, also downloaded from cBioPortal and UCSC Xena. As an independent validation dataset, we obtained data from the Glioma Longitudinal Analysis (GLASS) Consortium, which is a global collaboration dedicated to collecting and analyzing longitudinal genomic and molecular data from patients with glioma.^26,33^ Using cBioPortal, we downloaded clinical, gene expression, and mutation data for primary tumors in the Diffuse Glioma GLASS dataset (version 2022-05-31).^34,35^

#### 2.1.1. Clinical data

Both the TCGA and GLASS datasets contain clinical data per patient and per sample. As outlined in Figure 1, we defined TCGA patients with glioma recurrences as patients meeting one of the following criteria: (1) patients with recurrence samples in the dataset; (2) patients with a non-zero value for *days to tumor recurrence*; (3) patients with *new neoplasm event type* equal to ‘Recurrence’; or (4) patients with both a new tumor event (NTE; indicated by *NTE after initial treatment* equal to ‘Yes’, *disease-free status* equal to ‘Recurred/Progressed’, or a non-zero value for *days to NTE after initial treatment*) <u>and</u> successful treatment of the primary tumor (indicated by *primary therapy outcome success* or *treatment outcome first course* equal to ‘Complete Remission/Response’). For patients with recurrence, we defined time to recurrence (TTR) by *days to tumor recurrence* (otherwise *days to NTE after initial treatment* or *disease-free status months*). Unlike TCGA, the GLASS dataset contains surgical timelines. As such, we identified GLASS patients with recurrence based on the presence or absence of a recurrence surgery and defined TTR as the time elapsed between surgery for primary tumors and surgery for first recurrence tumors (based on *start date*). For both TCGA and GLASS, the median TTR was between 1 and 1.5 years; therefore, we labeled patients with less than 1 year to recurrence as *early* and patients with more than 1 year to recurrence as *late*. As LUNAR is designed to predict recurrence based on data available at the time of primary diagnosis, we removed any clinical features that encode information not available at the point of primary tumor treatment, such as overall survival. We also relabeled patients whose gliomas were classified using pre-2021 WHO guidelines (Supplementary Methods and Figure S1).

**Figure 1.**
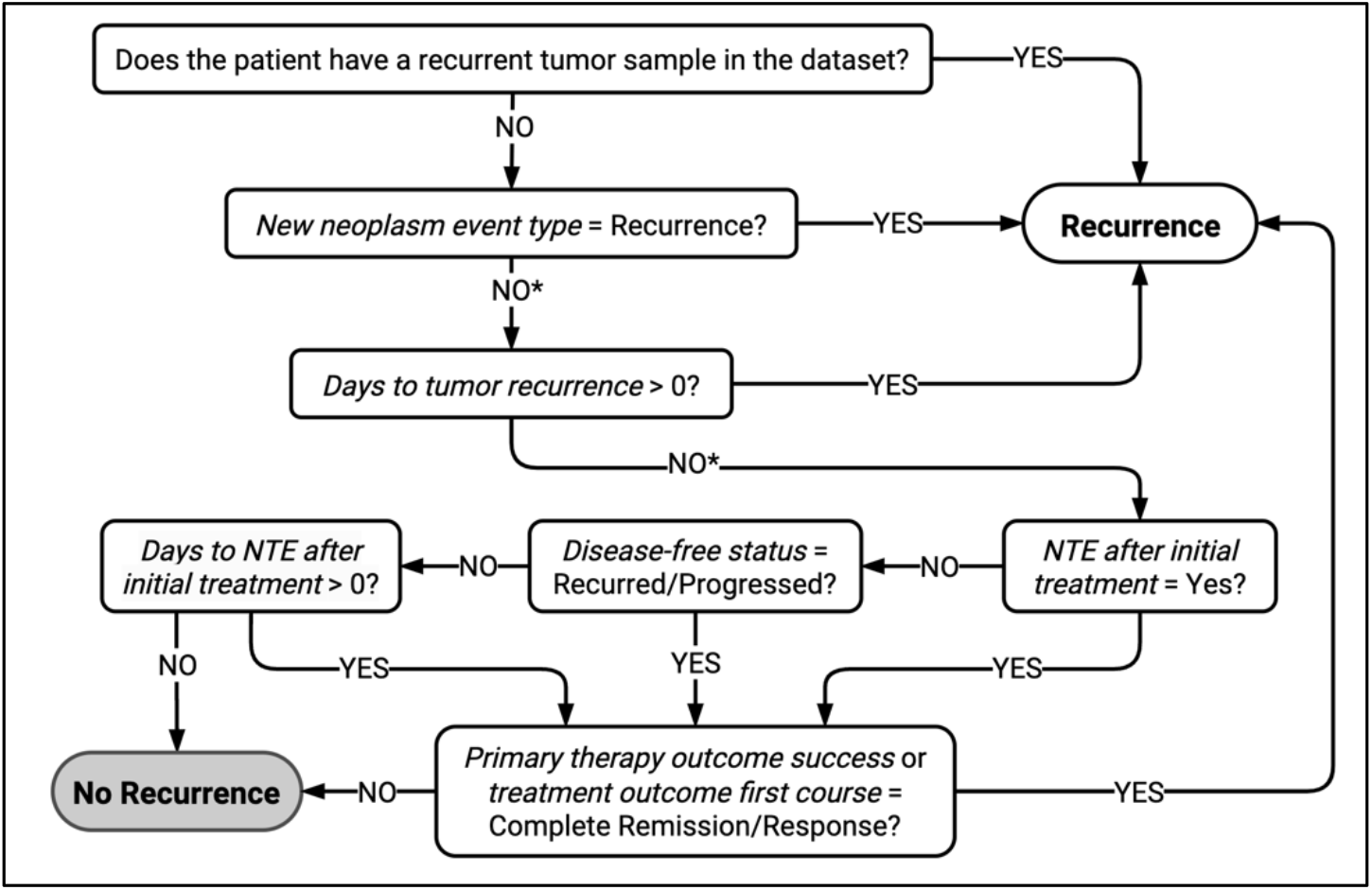
The workflow used to label TCGA patients as with or without glioma recurrence. *Days to tumor recurrence was only available for the individual GBM dataset; NTE = new tumor event. *Indicates the path also includes patients for which the given variable is not recorded.

#### 2.1.2. Genomic data

We downloaded gene expression data and Mutation Annotation Format files containing somatic mutations for all sequenced patients in the GLASS and TCGA datasets. For both datasets, the gene expression data were log2(x+1) transformed and mean-normalized per gene across their respective repositories prior to download. From the mutation data, we excluded silent mutations and calculated the number of mutations per gene per patient. The distribution of the variant counts based on the variant classification for TCGA and GLASS patients are available in Supplementary Figure S2 and Supplementary Figure S3, respectively. Variant classifications included translation start site, splice site, splice region, intron, nonsense, nonstop, missense, in-frame, frame shift, 5’ UTR, 3’ UTR, 5’ flank, and 3’ flank mutations.

### 2.2. Data preprocessing

To standardize gene names across genomic data and filter the genomic data to the most relevant genes, we downloaded all currently approved protein-coding genes located on the 22 autosomes from the HUGO Gene Nomenclature Committee BioMart Gene repository.^36^ We then narrowed the data to genes in the downloaded repository, mapping by Entrez ID or gene symbol. To avoid unintentional exclusion of genes due to changes in official symbols, we checked alias symbols and former symbols in addition to approved symbols for all genes in the dataset. The resulting genomic features for each patient were mutation counts per gene (where each feature represents the number of mutations for a single gene) and mRNA expression per gene (where each feature represents the expression for a single gene).

After applying the criteria mentioned above and restricting to patients with clinical, gene expression, and mutation data, we withheld 20% of patients from each dataset for final evaluation (excluded from all training, feature selection, hyperparameter tuning, and further preprocessing). Of the remaining data, we chose an 80/20 stratified training and validation split.

For each dataset, we used the training data to remove features with only one unique value and features with missing values. Both datasets demonstrated a high degree of collinearity among features. Thus, we selected all highly correlated feature pairs (Pearson pairwise correlation 0.95) and, for each pair, removed whichever feature had the larger number of high correlations among all features. This led to the removal of 7716 constant TCGA features (165 expression, 7551 mutation), 3915 constant GLASS features (9 expression, 3906 mutation), 1355 highly correlated TCGA features (118 expression, 1237 mutation), and 5403 highly correlated GLASS features (1 clinical, 164 expression, 5238 mutation). Lastly, we applied Scikit-learn RobustScaler^37^ to all mutation and expression features and continuous clinical features. After all preprocessing, the TCGA dataset contained 17,534 total features (13 clinical, 17,273 expression, and 248 mutation) while the GLASS dataset contained 25 143 total features (16 clinical, 17,826 expression, and 7301 mutation).

### 2.3. Final patient cohort

The TCGA dataset had 648 patients with clinical, gene expression, and mutation data meeting our criteria. Of those patients, 19 had recurrent samples in the dataset, 13 had an explicit recurrence recorded (via recurrence-specific variables, when available), and 125 had both a NTE and successful treatment of their primary tumors. Thus, 157 TCGA patients were included in the TCGA cohort (Table 1). Of the TCGA patients included, 41 (26.1%) had recurrence less than 1 year after initial treatment, and 116 (73.9%) had at least 1 year between initial treatment and recurrence. The majority of patients with grade IV glioma (73.7%) had early recurrences, versus 22.2% of patients with grade III and 16.7% of patients with grade II. IDH-mutant gliomas were dominant across recurrence groups (n=126, 80.3%), 38.1% (n=48) of which had 1p19q co-deletion. The three most frequently mutated genes in TCGA patients included *IDH1, TP53*, and *TTN*.

**Table 1.** Descriptive statistics for the TCGA cohort according to recurrence outcome and tumor grade.

| Tumor grade* | | Early recurrence or progression (<1 yr.) | | | | Late recurrence or progression ($\geq 1$ yr.) | | | |
| --- | --- | --- | --- | --- | --- | --- | --- | --- | --- |
|  |  | All (41) | II (11) | III (16) | IV (14) | All (116) | II (55) | III (56) | IV (5) |
| Sex [n] | Male | 23 | 6 | 8 | 9 | 62 | 26 | 33 | 3 |
|  | Female | 18 | 5 | 8 | 5 | 54 | 29 | 23 | 2 |
| Race<sup>a</sup> [n] | White | 37 | 10 | 16 | 11 | 109 | 49 | 55 | 5 |
|  | African American or Black | 2 | 1 | 0 | 1 | 2 | 2 | 0 | 0 |
|  | Asian | 1 | 0 | 0 | 1 | 1 | 0 | 1 | 0 |
| Hispanic or Latino<sup>a</sup> [n] | No | 38 | 10 | 16 | 12 | 103 | 44 | 54 | 5 |
|  | Yes | 1 | 1 | 0 | 0 | 6 | 6 | 0 | 0 |
| Tumor type* [n] | ASTR | 14 | 6 | 8 | 0 | 64 | 32 | 31 | 1 |
|  | GBM | 14 | 0 | 0 | 14 | 4 | 0 | 0 | 4 |
|  | ODG | 10 | 4 | 6 | 0 | 38 | 20 | 18 | 0 |
|  | ASTR - WT | 3 | 1 | 2 | 0 | 10 | 3 | 7 | 0 |
| IDH & 1p19q codeletion status* [n] | WT | 17 | 1 | 2 | 14 | 14 | 3 | 7 | 4 |
|  | Mut, non-codel | 14 | 6 | 8 | 0 | 64 | 32 | 31 | 1 |
|  | Mut, codel | 10 | 4 | 6 | 0 | 38 | 20 | 18 | 0 |
| Survival status* [n] | Alive | 26 | 10 | 14 | 2 | 101 | 51 | 50 | 0 |
|  | Deceased | 15 | 1 | 2 | 12 | 15 | 4 | 6 | 5 |
| Months survival* | Median | 10.4 | 6.4 | 9.0 | 14.3 | 28.8 | 28.5 | 29.2 | 29.0 |
|  | Q<sub>1</sub>, Q<sub>3</sub> | 6.0, 14.3 | 3.7, 10.7 | 5.4, 12.0 | 10.5, 17.8 | 18.0, 43.4 | 18.2, 44.3 | 18.5, 41.4 | 18.0, 47.6 |
| Days to recurrence | Median | 193 | 193 | 244.5 | 131.5 | 787.5 | 704 | 845.5 | 654 |
|  | Q<sub>1</sub>, Q<sub>3</sub> | 105.0, 291.0 | 113.5, 282.0 | 163.8, 320.0 | 79.8, 247.5 | 513.3, 1142.0 | 501.5, 1051.0 | 540.3, 1193.3 | 427.0, 797.0 |
| Age [yr]* | Median | 47 | 38 | 43.5 | 58.5 | 37.5 | 34 | 39 | 56 |
|  | Q<sub>1</sub>, Q<sub>3</sub> | 36.0, 60.0 | 30.0, 43.0 | 35.8, 60.3 | 53.8, 62.0 | 30.0, 48.0 | 29.0, 45.0 | 32.8, 50.3 | 49.0, 63.0 |

ASTR = astrocytoma; GBM = glioblastoma; ODG = oligodendroglioma; WT = wildtype; mut = mutant; codel = 1p19q codeletion; \* $P < .005$ .
<sup>a</sup>Race and ethnicity were unavailable for 5 and 9 patients, respectively.

Of the 329 patients with clinical data in the GLASS dataset, 281 patients had recurrent and primary tumors recorded. Of those patients, 146 had gene expression data, 257 had mutation data, and 121 had both expression and mutation data. Of the 121 patients included in the GLASS cohort (Table 2), 60 (49.6%) had recurrences less than 1 year after initial treatment, and 61 (50.4%) had at least 1 year between initial treatment and recurrence. Grade IV tumors were most common (n=92, 76.0%), while 16 patients (13.2%) and 13 patients (10.7%) had grade II and grade III tumors, respectively. Grade II and grade III tumors accounted for 8.3% of early recurrence cases and 39.3% of late recurrence cases. IDH-wildtype gliomas were dominant across recurrence groups. Of patients with IDH-mutant gliomas (n=25, 20.7%), 20% (n=5) had 1p19q co-deletion. The three most frequently mutated genes in the GLASS dataset included *RYR2, EYS*, and *TP53*.

**Table 2.** Descriptive statistics for the GLASS cohort according to recurrence outcome and tumor grade.

| Tumor grade* |  | Early recurrence or progression (< 1 yr.) |  |  |  | Late recurrence or progression (≥1 yr.) |  |  |  |
| --- | --- | --- | --- | --- | --- | --- | --- | --- | --- |
|  |  | All (60) | II (3) | III (2) | IV (55) | All (61) | II (13) | III (11) | IV (37) |
| Sex [n] | Male | 42 | 2 | 1 | 39 | 32 | 4 | 9 | 19 |
|  | Female | 18 | 1 | 1 | 16 | 29 | 9 | 2 | 18 |
| Tumor type* [n] | GBM | 54 | 0 | 0 | 54 | 35 | 0 | 0 | 35 |
|  | ASTR | 4 | 3 | 0 | 1 | 16 | 10 | 4 | 2 |
|  | ASTR - WT | 2 | 0 | 2 | 0 | 5 | 0 | 5 | 0 |
|  | ODG | 0 | 0 | 0 | 0 | 5 | 3 | 2 | 0 |
| IDH & 1p19q codeletion status* [n] | WT | 56 | 0 | 2 | 54 | 40 | 0 | 5 | 35 |
|  | Mut, non-codel | 4 | 3 | 0 | 1 | 16 | 10 | 4 | 2 |
|  | Mut, codel | 0 | 0 | 0 | 0 | 5 | 3 | 2 | 0 |
| Survival status [n] | Alive | 7 | 2 | 1 | 4 | 7 | 4 | 1 | 2 |
|  | Deceased | 53 | 1 | 1 | 51 | 54 | 9 | 10 | 35 |
| Months survival* | Median | 17 | 40 | 31.5 | 17 | 39 | 64 | 46 | 25 |
|  | Q<sub>1</sub>, Q<sub>3</sub> | 13.0, 22.0 | 31.5, 80.5 | 22.3, 40.8 | 13.0, 21.0 | 23.0, 56.0 | 50.0, 94.0 | 33.5, 77.0 | 21.0, 43.0 |
| Days to recurrence | Median | 243 | 274 | 289 | 213 | 669 | 1156 | 1125 | 517 |
|  | Q<sub>1</sub>, Q<sub>3</sub> | 152.0, 304.0 | 228.5, 304.5 | 266.0, 312.0 | 152.0, 304.0 | 487.0, 1156.0 | 760.0, 1703.0 | 654.0, 1444.5 | 426.0, 821.0 |
| Age [yr]* | Median | 56.5 | 43 | 47.5 | 57 | 49 | 37 | 35 | 54 |
|  | Q<sub>1</sub>, Q<sub>3</sub> | 47.0, 64.0 | 36.0, 43.0 | 45.8, 49.3 | 48.5, 64.5 | 37.0, 60.0 | 32.0, 42.0 | 24.5, 45.5 | 48.0, 63.0 |
| Stromal score | Median | -488.8 | -940.6 | -1014.7 | -477.8 | -701.4 | -1446.5 | -993.4 | -412.9 |
|  | Q<sub>1</sub>, Q<sub>3</sub> | -768.5, -111.8 | -1254.2, -852.9 | -1068.8, -960.6 | -657.7, -77.3 | -1109.1, -299.8 | -1527.0, -1068.8 | -1131.3, -887.3 | -830.9, 24.6 |
| Immune score* | Median | 584.8 | 249.2 | 95.9 | 613.3 | 201.7 | -431.4 | -287.5 | 468.9 |
|  | Q<sub>1</sub>, Q<sub>3</sub> | 176.2, 926.5 | -186.9, 290.4 | 66.6, 125.3 | 260.3, 957.8 | -380.3, 749.8 | -720.6, -248.3 | -478.1, 257.7 | 142.7, 892.0 |
| ESTIMATE score | Median | 155.1 | -691.4 | -918.8 | 268.0 | -527.2 | -1792.9 | -1319.4 | 317.7 |
|  | Q<sub>1</sub>, Q<sub>3</sub> | -505.7, 733.1 | -1441.1, -562.5 | -943.5, -894.1 | -382.8, 775.4 | -1540.5, 464.5 | -2232.2, -1306.6 | -1592.4, -629.6 | -527.2, 879.1 |
| Purity | Median | 0.81 | 0.88 | 0.89 | 0.8 | 0.86 | 0.94 | 0.92 | 0.8 |
|  | Q<sub>1</sub>, Q<sub>3</sub> | 0.76, 0.86 | 0.87, 0.92 | 0.89, 0.89 | 0.75, 0.85 | 0.78, 0.93 | 0.92, 0.96 | 0.87, 0.93 | 0.74, 0.86 |

ASTR = astrocytoma; GBM = glioblastoma; ODG = oligodendroglioma; WT = wildtype; mut = mutant; codel = 1p19q codeletion. \*P<.005.

### 2.4. LUNAR

We developed g**L**ioma rec**U**rre**N**ce **A**ttention-based classifie**R** (LUNAR) using clinical, expression, and mutation data from patients with glioma to predict early and late recurrence. The PyTorch (version 3.2.1)^38^ framework for LUNAR is outlined in Figure 2. First, each data type (clinical, expression, and mutation) is individually passed through modality-specific neural networks (NNs) with 3 fully connected layers. The outputs of each NN enter a four-headed self-attention layer. Self-attention is a variant of attention in which each element in an input weighs its relevance to all other elements, including itself, allowing the model to learn internal interactions among the input and determine which aspects are most important for making predictions. In the context of LUNAR, the modality-specific self-attention layers capture intramodal relationships (i.e., relationships between features of the same modality) and identify which features to prioritize for each of the three input types.

**Figure 2.**
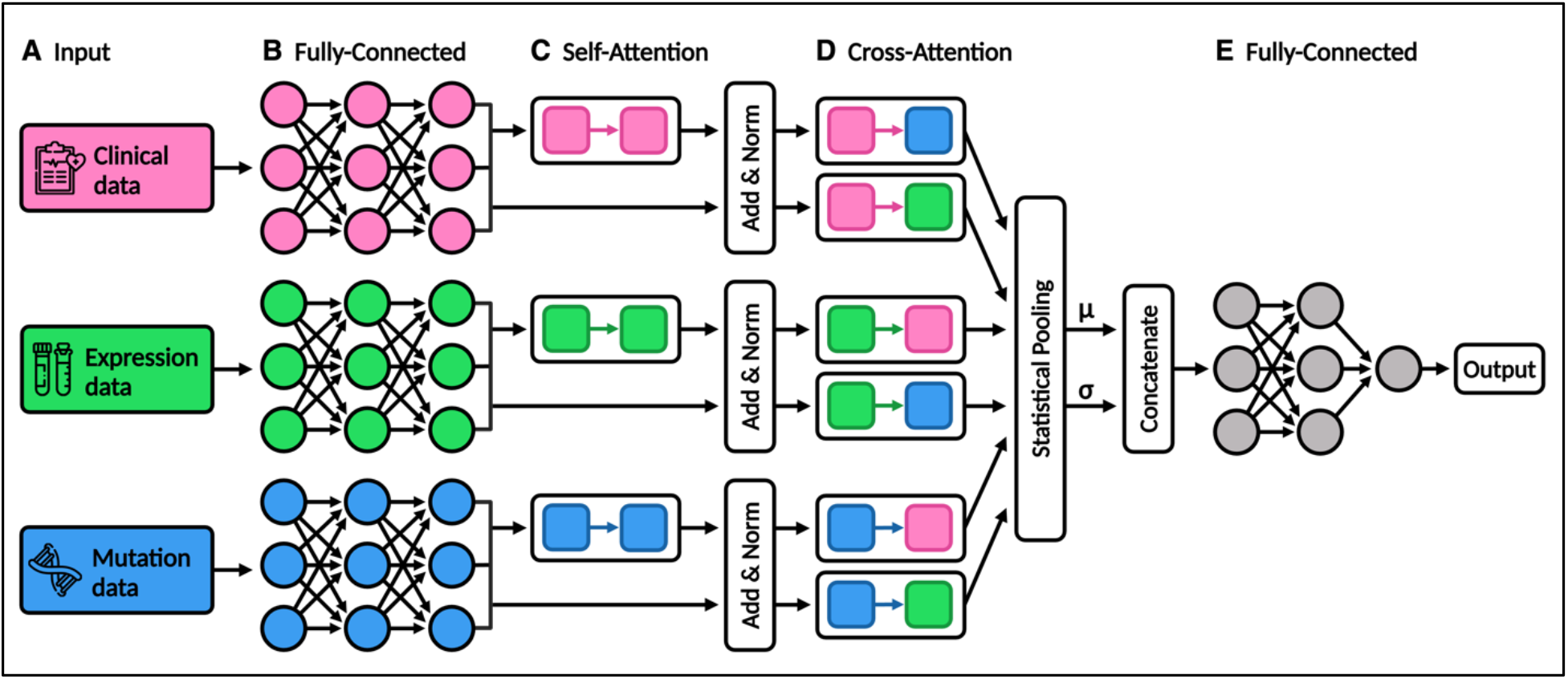
The LUNAR framework: (A) LUNAR accepts tabular clinical, expression, and mutation count data. (B) Inputs are passed to modality-specific fully connected (FC) neural networks. (C) Outputs are passed to a multi-head self-attention layer. The outputs of the NNs and self-attention layers are residually connected and normalized. (D) Outputs for every modality pairing are passed to bi-directional cross-attention layers, after which they are statistically pooled, concatenated, and passed to (E) two final FC layers and a Sigmoid output layer.

Next, outputs from the fully connected NNs and the single-modality self-attention layers are added together (residually connected) and normalized. By passing information from earlier layers and normalizing the outputs across each modality’s features, this technique (Add & Norm)^39^ serves to both stabilize the training process and restore any potentially diminishing gradients. To capture intermodal or cross-modal relationships (i.e., relationships between features of different modalities), the normalized outputs for each possible pairing of modalities are passed to cross-modality attention (cross-attention) layers. In contrast to the unidirectional self-attention layers described above, our cross-attention layers are bidirectional and comprised of two unidirectional attention calculations.^21^ For example, the cross-attention between clinical output matrix and mutation output matrix is the result of concatenation(cross-attention(**C** → **M**), cross-attention(**M** → **C**)). The outputs of the cross-attention layers are then statistically pooled, concatenated, and processed through two final fully connected layers. By using both self-attention and cross-attention, the model first learned the most important features within a given modality and then the most important features between modalities.

#### 2.3.3. Feature selection and hyperparameter tuning

To optimize the LUNAR models, we first created different sets of selected expression and mutation features using ANOVA F-score selection and selection via model importance for 4 classifiers: 1) L1-penalized logistic regression; 2) linear support vector classifier (SVC); 3) random forest; 4) eXtreme Gradient Boosting (XGBoost).^37,40^ Selector parameters were determined using a randomized 3-fold cross-validation hyperparameter search. We then tuned the sets of selected expression and mutation features in combination with model hyperparameters, including modality-specific network architecture (network depth, layer dimensions, dropout rate), self- and cross-attention dropout rates, activation functions, learning rate, and optimization algorithm, to find the optimal combination of features and parameters. Model parameters and specifications are detailed in Supplementary Table S1. Clinical data features did not undergo feature selection due to the comparatively low number of clinical features in the TCGA and GLASS datasets. For the LUNAR model fine-tuned on the TCGA dataset (LUNAR TCGA), model performance was best when using expression features selected via XGBoost (24 features) and mutation features selected via ANOVA F-score (24 features). LUNAR GLASS performed best when using expression features selected via random forest (1994 features) and mutation features selected via ANOVA F-score (730 features). Both TCGA and GLASS had clinical features representing patient age, tumor classification, tumor grade, IDH mutation and 1p19q codeletion status, and patient sex. GLASS clinical features also included stromal score, immune score, and ESTIMATE score. Glioma tissues contain a multitude of glioma-associated non-tumor cells in their microenvironment that dilute the purity of glioma cells and play important roles in glioma development, specifically infiltrating immune cells and stromal cells.^41^ Therefore, Yoshihara et al.^42^ designed the ESTIMATE (Estimation of Stromal and Immune Cells in Malignant Tumours using Expression Data) algorithm, which calculates a stromal score (estimated presence of stromal cells), immune score (estimated infiltration of immune cells), and composite ESTIMATE score (estimated non-tumor cell content), to quantify overall tumor purity in tumor tissues.

### 2.5. Model evaluation and training

To assess model performance, we benchmarked our models against four traditional ML classifiers used in cancer recurrence studies by González-Castro et al.^15^ and Zuo et al.^16^ These models include SVC, logistic regression (L2-penalized), decision tree, and Gaussian naive Bayes (GaussianNB).^37^ All models compared used the same data and selected features. For each model, we calculated AUROC, area under the precision-recall curve (AUPRC), accuracy, balanced accuracy, precision, recall (sensitivity), specificity, and F1-score. To evaluate the influence of attention mechanisms on LUNAR performance, we also tested two additional LUNAR models: LUNAR without cross-attention and LUNAR without cross- and self-attention.

All computational work for this study was performed on a high-performance computing cluster, Ocean State Center for Advanced Resources (Oscar),^43^ at Brown University Center for Computation and Visualization. We used 1 GPU (NVIDIA GeForce RTX 3090) for model training and 4 GPUs for parallelized fine-tuning.

### 2.6. Explainability

To evaluate the impact of each feature on LUNAR’s predictions, we utilized SHapley Additive exPlanations (SHAP) DeepExplainer.^44,45^ In the SHAP method, each feature’s relative contribution to the model prediction is quantified as a SHAP value, representing the average marginal contribution across all possible feature combinations. Positive SHAP values indicate a feature has a positive impact on model predictions, pushing the model towards an early recurrence prediction, while those with negative values have a negative influence and push the model towards a late recurrence prediction. Feature importance is calculated by averaging a given feature’s absolute SHAP values across the dataset.

## 3. Results

### 3.1. Model Performance

Both LUNAR TCGA and LUNAR GLASS outperformed all benchmarked models in each metric tested. LUNAR TCGA achieved an AUROC of 90.63%, AUPRC of 75.86%, accuracy of 87.5%, precision of 75.0%, recall of 75.0% (tied with its no-attention counterpart and linear SVC), and specificity of 91.67% (tied with its self-attention-only counterpart). LUNAR GLASS achieved an AUROC of 89.1%, AUPRC of 89.4%, accuracy of 84.0%, precision and recall of 83.33%, and specificity of 84.62%. LUNAR GLASS self-attention-only was the second highest performer in all metrics compared to the GLASS baselines, while LUNAR TCGA self-attention-only was the second highest performer in AUPRC, accuracy, and precision. Receiver operating characteristic (ROC) and precision-recall (PR) curves are shown in Figure 3, while full performance metrics for all classifiers are available in Supplementary Table S2 (TCGA) and Table S3 (GLASS).

**Figure 3.**
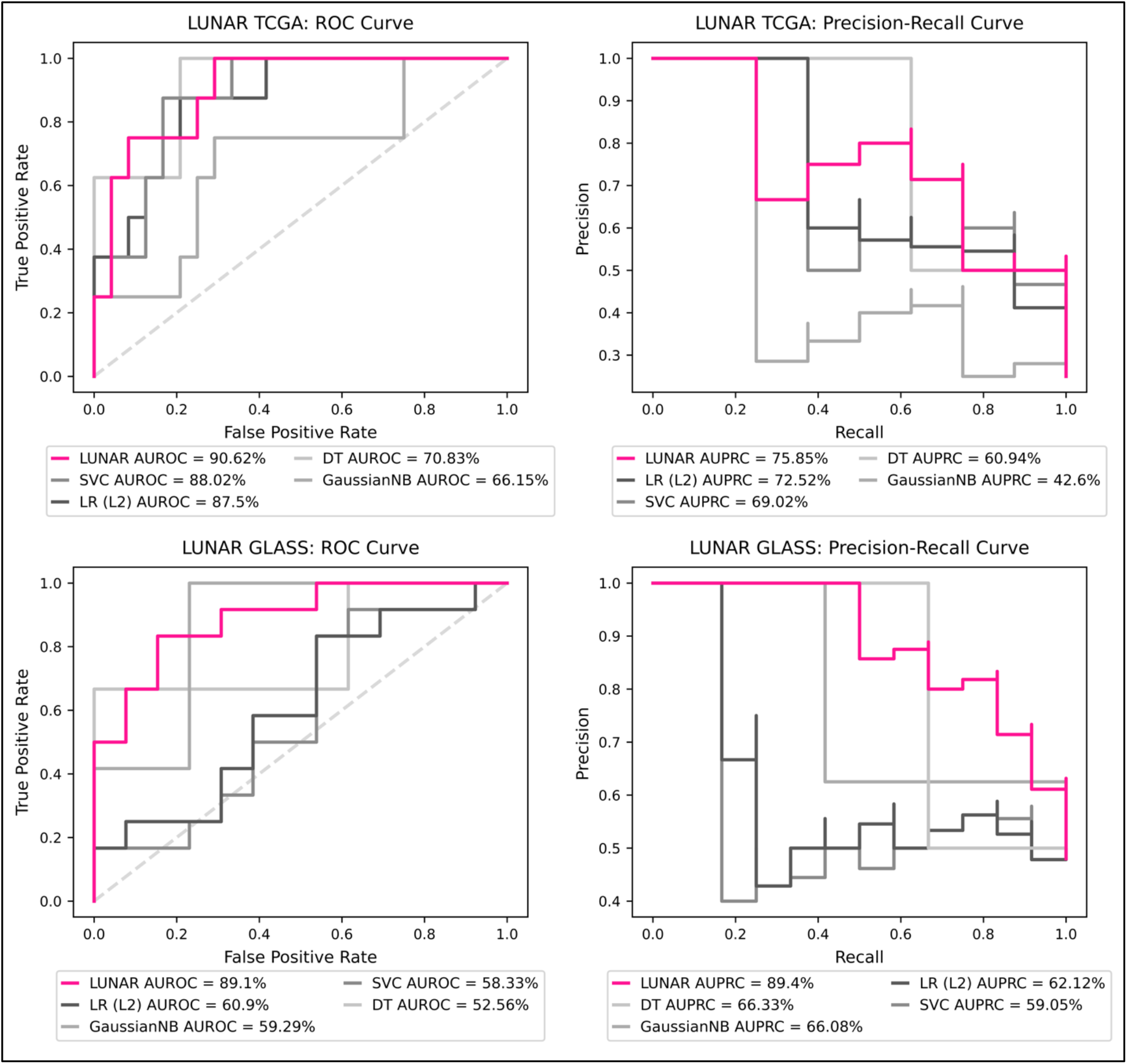
ROC (left) and PR (right) curves comparing LUNAR and the four traditional machine learning models’ performance on the TCGA (top) and GLASS (bottom) datasets. LR = logistic regression; SVC = support vector classifier; DT = decision tree; NB = naive Bayes.

### 3.2. Feature importance

SHAP summary plots for the 20 most important features to LUNAR TCGA and LUNAR GLASS are presented in Figure 4 and Figure S4, respectively. For both models, clinical features had the highest absolute mean importance per feature, while expression features had the lowest absolute mean importance per feature. Features that appeared in the top 50 features for both datasets include patient age, sex, IDH mutation status, tumor grade, and tumor classification. The top 30 most important features to LUNAR TCGA are 13.3% (n=4) clinical, 53.3% (n=16) expression, and 33.3% (n=10) mutation. The top 30 most important features to LUNAR GLASS are 36.7% (n=11) clinical, 16.7% (n=5) expression, and 46.7% (n=14) mutation.

**Figure 4.**
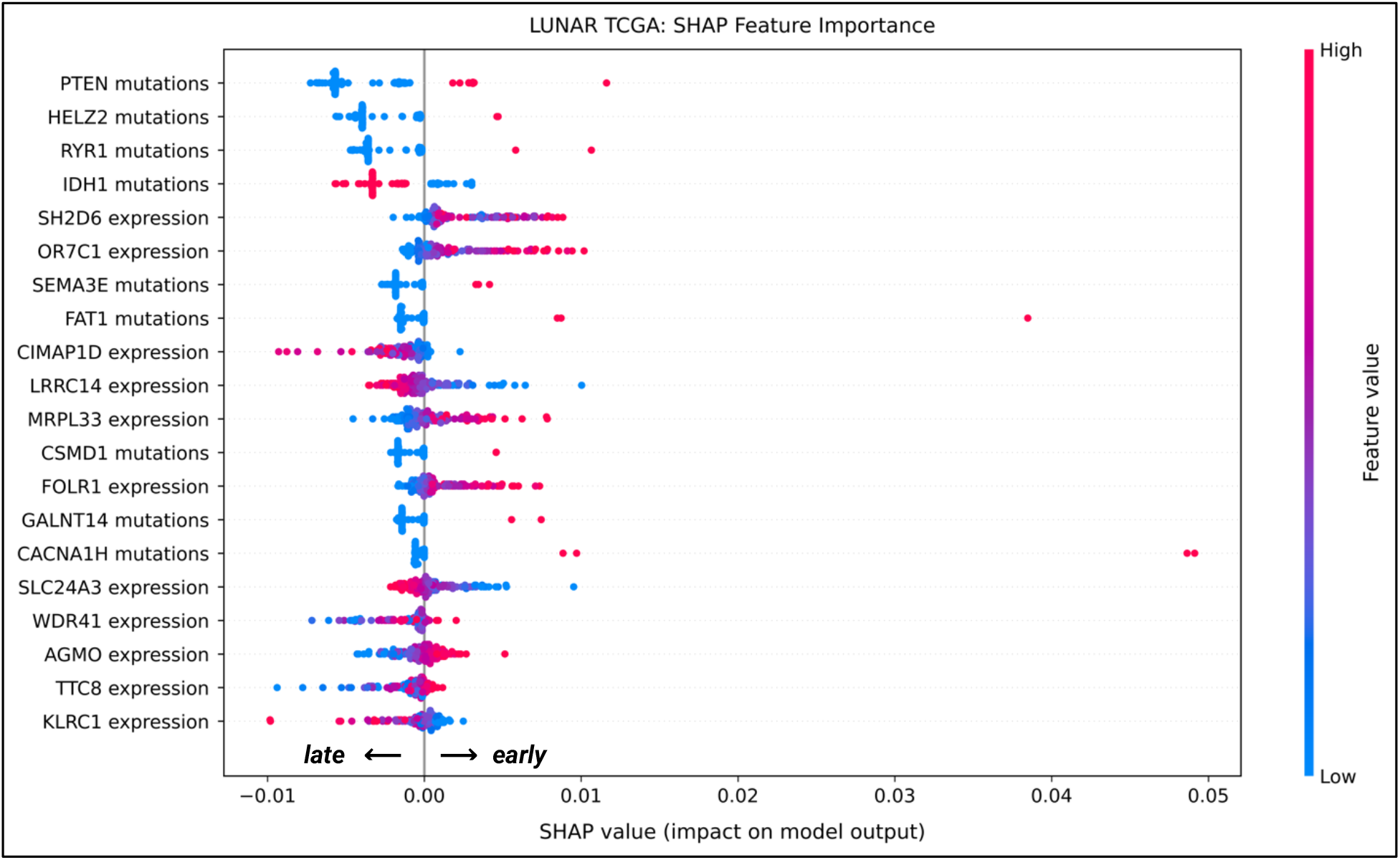
The 20 most influential features for LUNAR TCGA according to DeepExplainer. Positive SHAP values (dots to the right) indicate an increase in the model’s prediction (towards an early prediction), while negative values (dots to the left) indicate a decrease (towards a late prediction). Note that for categorical features, red = YES and blue = NO.

## 4. Discussion

LUNAR, an attention-based DL model outperformed traditional ML models using both TCGA and GLASS datasets in predicting glioma recurrence. Clinical features were found to have the greatest importance in LUNAR recurrence predictions in both the GLASS and TCGA datasets, followed by mutation counts and mRNA expression. According to SHAP DeepExplainer, LUNAR TCGA and LUNAR GLASS both tended to output early recurrence predictions for patients who had IDH-wildtype tumors and late recurrence predictions for patients who had IDH-mutant tumors. LUNAR TCGA was further influenced towards late recurrence when IDH-mutant tumors also exhibited 1p19q co-deletion. These patterns align with established literature, which has shown mutations in *IDH1/2* and 1p19q co-deletion are associated with increased response to treatment and longer overall survival, and are more common in LGG.^1,46–49^ This may also explain why oligodendrogliomas, which require 1p19q co-deletion, also pushed LUNAR TCGA towards late recurrence predictions.^50^

LUNAR GLASS was also more likely to predict early recurrence for patients with high immune scores (increased infiltration of immune cells) and low ESTIMATE scores (decreased tumor purity). Studies have shown that stromal cells and interactions between glioma and immune cells can establish a dynamic tumor microenvironment (TME) that facilitates tumor proliferation and invasion, increases angiogenesis, encourages treatment resistance, and contributes to an immunosuppressive microenvironment.^51–55^ Zhang et al.^56^ found in four glioma cohorts that gliomas with low purity and high stromal and immune scores were more aggressive and conferred shorter overall survival than gliomas with higher purity and lower stromal and immune scores.

Multiple mutation and expression features of high SHAP importance similarly have established relationships with glioma. For example, SHAP importance indicates LUNAR TCGA was influenced towards early recurrence predictions in the presence of *PTEN, SEMA3E, FAT1*, and *SETD2* mutations. *PTEN* encodes a tumor suppressing dual-specificity phosphate involved in negative regulation of protein phosphorylation and protein dephosphorylation that suppresses tumor growth by antagonizing the PI3K-AKT/PKB signaling pathway.^57^ Multiple studies have demonstrated *PTEN* mutations in glioma are associated with significantly shorter overall survival and therapeutic resistance.^58,59^ *SEMA3E* encodes a class-3 semaphorin and axon guidance factor essential to hypothalamic neuron development.^60^ In a study on class-3 semaphorins, Sabag et al.^61^ found glioblastoma cells expressing *SEMA3E* significantly inhibited tumor development following implantation subcutaneously and implantation in the cortex of mouse brains (separately), and more than doubled mouse survival time. *FAT1* encodes a transmembrane protein that can act as both a tumor suppressor gene and an oncogene in various cancers.^62,63^ Morris et al.^64^ found *FAT1* knockdown in glioma cells led to consistent enrichment of the Wnt/β-catenin signaling pathway, while GBM cell lines that expressed non-mutated FAT1 suppressed rate of cell growth. *SETD2* encodes an H3K36 trimethyltransferase involved in angiogenesis, vasculogenesis, and neural tube closure. Viaene et al.^65^ and Fontebasso et al.^66^ demonstrated significant associations between *SETD2* mutations and both pediatric and adult HGGs. LUNAR TCGA early recurrence predictions were also positively influenced by high *FOLR1* expression and negatively influenced by high *KLRC1* expression. *FOLR1* encodes a cell-surface glycoprotein responsible for coordinating the transport of folate and its derivants.^67^ *FOLR1* has shown significant upregulation in multiple cancers, particularly glioma, which has made it an attractive target for intervention.^68,69^ *KLRC1* encodes a transmembrane protein preferentially expressed in natural killer cells, and in the Chinese Glioma Genome Atlas, Sun et al.^70^ found high tumor expression of *KLRC1* was associated with improved prognosis.

While the results of our study are promising, there are several limitations and opportunities for improvement in future works. First, while the publicly available TCGA and GLASS datasets are highly valuable community resources, our sample size was relatively small after restricting to patients with recurrence events and all three data types. As such, the importance of rigorous and comprehensive clinical annotations cannot be overstated.^71^ Second, our primary dataset had class imbalance and limited demographic diversity, with 73.9% of patients labeled as late recurrence and 93.0% of patients identified as white. To compensate for class imbalance and minimize the possibility of overfitting to site- or population-specific data points, we validated our approach on a separate non-overlapping dataset (GLASS) with significantly less imbalance (50.8% labeled as late recurrence). However, future validation on additional datasets with a more diverse patient population will be essential to determining model generalizability. Despite these limitations, our preliminary results suggest the usability of DL-based predictive models for cancer recurrence prediction.

## 5. Conclusion

The TCGA and GLASS data repositories have become an invaluable resource for genomic, epigenomic, transcriptomic, and proteomic research and have significantly advanced our understanding of glioma. Our findings underscore the importance of external validation in DL- and ML-based cancer genomics research. LUNAR outperformed traditional ML models using both TCGA and GLASS datasets, both with and without cross-attention layers, demonstrating the capacity of DL for meaningful pattern recognition in highly dimensional genomic datasets.

## Data Availability

Expression, mutation, and clinical data for the GLASS dataset were downloaded from cBioPortal ([GLASS link](https://www.cbioportal.org/study/summary?id=difg_glass)). Clinical data for the TCGA datasets were downloaded from cBioPortal ([LGG link](https://www.cbioportal.org/study/summary?id=lgg_tcga), [GBM link](https://www.cbioportal.org/study/summary?id=gbm_tcga), [GBMLGG link](https://www.cbioportal.org/study/summary?id=lgggbm_tcga_pub)) and Xena (LGG link, GBM link, GBMLGG link). Mutation data and expression data for the GBMLGG dataset were downloaded from cBioPortal and Xena, respectively.

## Acknowledgements

We thank Brown University’s Center for Computation and Visualization for the computational resources, and Dr. J. Nicholas Fisk for their valuable feedback on this work.

