## Supplementary materials for "LUNAR: A Deep Learning Model to Predict Glioma Recurrence Using Integrated Genomic and Clinical Data"

#### Clinical Feature Relabeling Based on Updated WHO Guidelines

In 2021, the 5th edition of the World Health Organization (WHO) Classification of CNS Tumors (CNS5) introduced significant changes to glioma classification.[1] The 2016 classification previously categorized diffuse gliomas as astrocytic tumors, oligodendroglial tumors, and oligoastrocytic tumors. Astrocytic tumors included diffuse astrocytoma (IDH-mutant or -wildtype), anaplastic astrocytoma (IDH-mutant or -wildtype), primary glioblastoma (IDH-wildtype grade IV astrocytomas), and secondary glioblastoma (IDH-mutant grade IV astrocytomas). Oligodendroglial tumors, defined as IDH-mutant tumors with combined deletion of both the short arm of chromosome 1 and the long arm of chromosome 19 (1p19q), included oligodendroglioma and anaplastic oligodendroglioma.[2] These guidelines also included criteria for oligoastrocytomas and anaplastic oligoastrocytomas. As of CNS5, IDH-mutant tumors encompass astrocytomas (in the absence of 1p19q codeletion, or if 1p19q testing is unavailable, loss of *ATRX* or *TP53* mutations) and oligodendrogliomas (in the presence of 1p19q codeletion). CNS5 also stipulates IDH-wildtype as a criterion for glioblastoma diagnosis, relabeling IDH-mutant glioblastomas without 1p19q codeletion as astrocytomas.

Both the GLASS and TCGA datasets contain patients with pre-CNS5 tumor classifications. To relabel patients, we applied the guidelines using IDH mutation and 1p19q codeletion status, as shown in Supplementary Figure S1. While IDH-wildtype astrocytomas without molecular features of glioblastoma are rare, assessing the validity of such a diagnosis would require further analyses outside the scope of this study. Therefore, IDH-mutant gliomas labeled as astrocytomas were labeled as 'astrocytoma wildtype' to distinguish them from the relabeled astrocytomas. CNS5 also added molecular criteria for tumor grading and switched from roman numerals to Arabic numerals. However, regarding each dataset's gliomas to WHO 1-4 requires analyses outside the scope of this study. Patient sex and *IDH* mutation status were binarily encoded, while tumor grade and tumor classification were categorically encoded as multiple features.

#### Attention mechanisms

*Attention mechanisms* are functions in prediction frameworks that dynamically assign varying levels of importance to different subsets of input, highlighting those that are most relevant to decision-making.[3,4] By attending to subsets, attention is well-suited for controlling interactions between and extracting information from high dimensional input spaces. Input is provided to attention layers as queries and keys of hidden dimension $d_k$ and values of hidden dimension $d_v$, packaged into matrices $\mathbf{Q}$, $\mathbf{K}$, and $\mathbf{V}$. Attention weights are then calculated using scaled dot-product attention.[4]

$$\text{Attention}(\mathbf{Q}, \mathbf{K}, \mathbf{V}) = \text{softmax}\left(\frac{\mathbf{Q}\mathbf{K}^T}{\sqrt{d_k}}\right)\mathbf{V}$$

*Multi-head attention* improves upon the above by linearly projecting $\mathbf{Q}$, $\mathbf{K}$, and $\mathbf{V}$ multiple times into $h$ different sub-queries, sub-keys, and sub-values (for $h$ heads) and computing scaled dot-product attention for each projection in parallel.[4] The independent outputs are then concatenated and projected into the expected dimensions. As a result, models can jointly attend to information from different representation subspaces at different positions. Note that all references to attention from this point forward refer to multi-head attention.

$$\text{MultiHead}(\mathbf{Q}, \mathbf{K}, \mathbf{V}) = \text{concat}(\text{head}_1, ..., \text{head}_h)\mathbf{W}^O \ni \text{head}_i = \text{Attention}(\mathbf{Q}\mathbf{W}_i^Q, \mathbf{K}\mathbf{W}_i^K, \mathbf{V}\mathbf{W}_i^V)$$

where the projections are parameter matrices $\mathbf{W}_i^Q$, $\mathbf{W}_i^K$, $\mathbf{W}_i^V$ and $\mathbf{W}^O$.

In *self-attention*, queries, keys, and values are equal, enabling the attention mechanism to learn interactions among inputs. In *cross-attention* or cross-modality attention, the source modality serves as $\mathbf{K}$ and $\mathbf{V}$, and the target modality serves as $\mathbf{Q}$. In contrast to the unidirectional intramodality attention described above, cross-attention is bidirectional and comprised of two unidirectional attention calculations.[5] In other words, we calculate the cross-attention between two modalities, A and B, as:

$$\text{Cross-Att}_{\mathbf{A},\mathbf{B}} = \text{concat}(\text{MultiHead}(\mathbf{A},\mathbf{B},\mathbf{B}),\ \text{MultiHead}(\mathbf{B},\mathbf{A},\mathbf{A}))$$

### Figures and tables

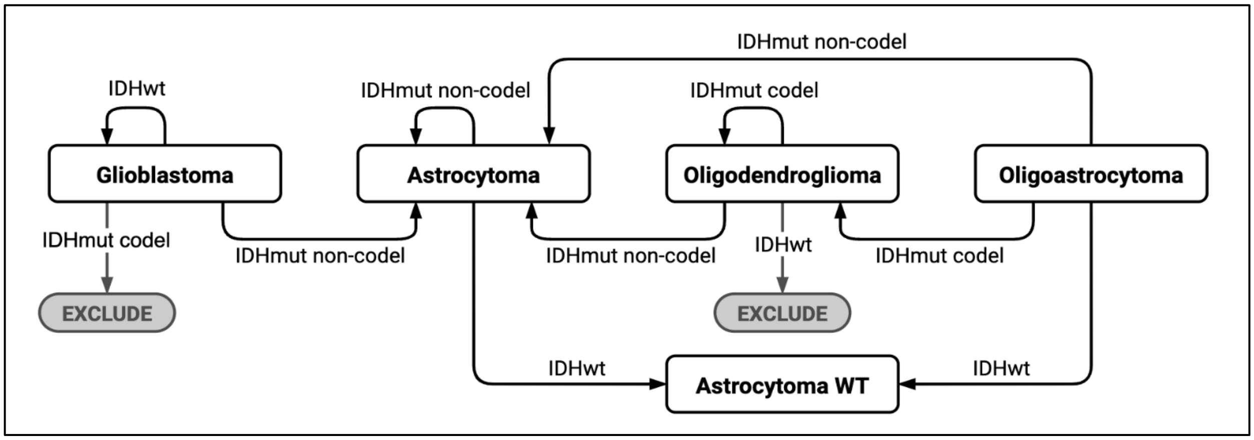


**Figure S1.** Algorithm used to relabel gliomas in accordance with the 2021 WHO Classification of CNS Tumors (CNS5). Determining the validity of IDH-wildtype astrocytoma designations fell outside the scope of this research, however we distinguished these tumors from IDH-mutant astrocytomas using the label 'Astrocytoma WT'; wt = wildtype; mut = mutant; codel = 1p19q co-deletion.

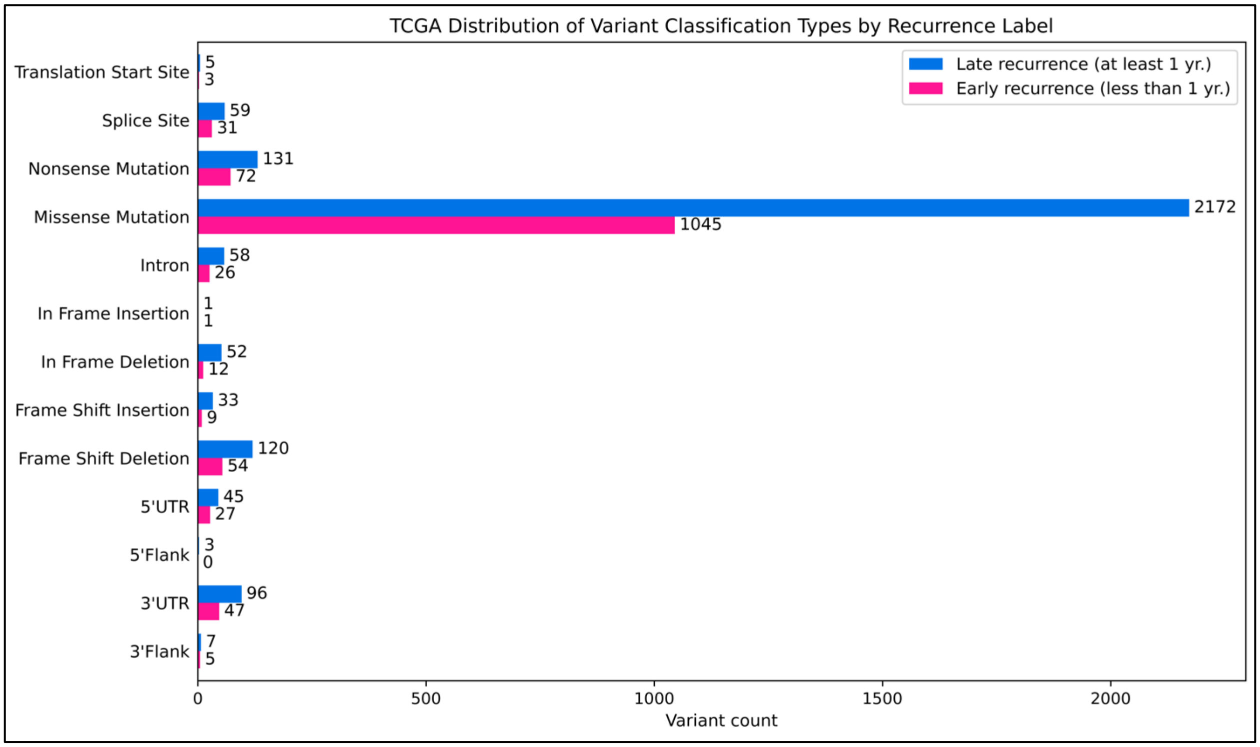


**Figure S2.** Distribution of mutations present in the TCGA cohort of primary gliomas.

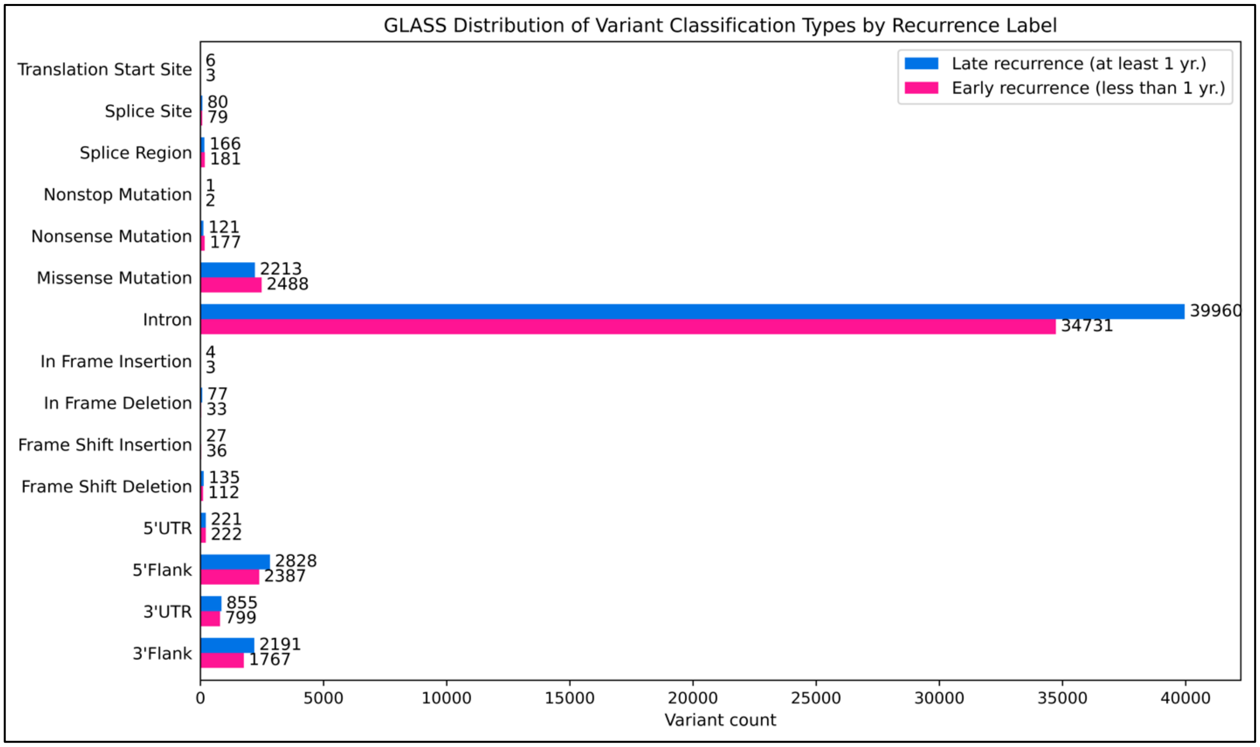


**Figure S3.** Distribution of mutations present in the GLASS cohort of primary gliomas.

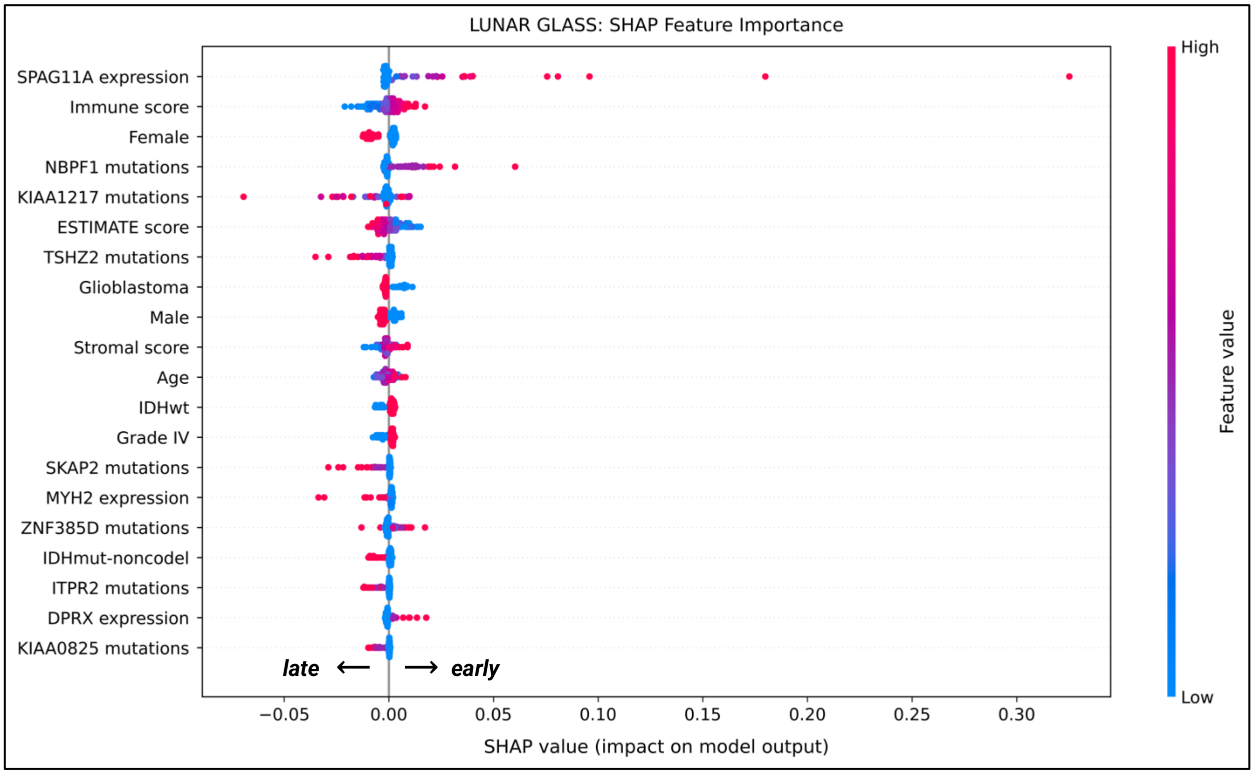


**Figure S4.** The 20 most influential features for LUNAR GLASS according to DeepExplainer. Positive SHAP values (dots to the right) indicate an increase in the model's prediction (towards an *early* prediction), while negative values (dots to the left) indicate a decrease (towards a *late* prediction). Note that for categorical features, red = YES and blue = NO.

**Table S1.** Model parameters, model configuration, and feature selection.

| | | TCGA | GLASS |
|---|---|---|---|
| **Clinical initial NN** | Neurons per hidden layer | 128, 128, 128 | 128, 128, 128 |
| | Dropout rate per hidden layer | 0.5, 0, 0.25 | 0, 0, 0.25 |
| | Activation function | ReLU | Leaky ReLU |
| **Expression initial NN** | Neurons per hidden layer | 256, 128, 128 | 256, 128, 128 |
| | Dropout rates per hidden layer | 0, 0.5, 0.25 | 0, 0, 0 |
| | Activation function | Leaky ReLU | Leaky ReLU |
| **Mutation initial NN** | Neurons per hidden layer | 512, 256, 128 | 256, 128, 128 |
| | Dropout rate per hidden layer | 0.5, 0.5, 0.25 | 0, 0, 0 |
| | Activation function | Leaky ReLU | Leaky ReLU |
| **Combined NN** | Neurons per hidden layer | 128 | 32 |
| | Activation function | Leaky ReLU | ReLU |
| **Feature selection** | Expression method (n selected) | XGBoost (24) | Random forest (1994) |
| | Mutation method (n selected) | Anova F-score (24) | Anova F-score (730) |
| **Configuration** | Optimizer | Adam | AdamW |
| | Learning rate | 0.001 | 0.0001 |
| | Weight decay | 0.00001 | 0.00001 |
| | Dataset sampling method | Weighted | Random |

**Table S2.** Performance metrics for LUNAR TCGA and the ML baselines.

| | AUROC | AUPRC | Accuracy | Balanced accuracy | Precision | Recall | Specificity | F1 |
|---|---|---|---|---|---|---|---|---|
| **LUNAR TCGA** | **90.63%** | **75.85%** | **87.50%** | **83.33%** | **75.0%** | **75.0%** | **91.67%** | **75.0%** |
| **Linear SVC** | 88.02% | 69.02% | 81.25% | 79.17% | 60.0% | **75.0%** | 83.33% | 66.67% |
| **L2 Logistic Regression** | 87.50% | 72.52% | 81.25% | 75.0% | 62.50% | 62.5% | 87.50% | 62.50% |
| **LUNAR TCGA (self-attention only)** | 85.94% | 74.24% | 84.38% | 77.08% | 71.43% | 62.5% | **91.67%** | 66.67% |
| **LUNAR TCGA (no attention)** | 82.29% | 63.40% | 78.13% | 77.08% | 54.55% | **75.0%** | 79.17% | 63.16% |
| **Decision Tree** | 70.83% | 60.94% | 75.0% | 70.83% | 50.0% | 62.5% | 79.17% | 55.56% |
| **Gaussian NB** | 66.15% | 42.60% | 65.63% | 52.08% | 28.57% | 25.0% | 79.17% | 26.67% |

AUROC = area under the receiver operating characteristic curve; AUPRC = area under the precision-recall curve; SVC = support vector classifier; NB = naive Bayes.

**Table S3.** Performance metrics for LUNAR GLASS and the ML baselines.

| | AUROC | AUPRC | Accuracy | Balanced accuracy | Precision | Recall | Specificity | F1 |
|---|---|---|---|---|---|---|---|---|
| **LUNAR GLASS** | **89.10%** | **89.40%** | **84.0%** | **83.97%** | **83.33%** | **83.33%** | **84.62%** | **83.33%** |
| **LUNAR GLASS (self-attention only)** | 78.85% | 81.52% | 72.0% | 72.44% | 66.67% | 83.33% | 61.54% | 74.07% |
| **LUNAR TCGA (no attention)** | 75.64% | 77.80% | 72.0% | 72.12% | 69.23% | 75.0% | 69.23% | 72.0% |
| **L2 Logistic Regression** | 60.90% | 62.12% | 52.0% | 52.24% | 50.0% | 58.33% | 46.15% | 53.85% |
| **Gaussian NB** | 59.30% | 66.08% | 60.0% | 59.30% | 62.50% | 41.67% | 76.92% | 50.0% |
| **Linear SVC** | 58.33% | 59.05% | 48.0% | 48.08% | 46.15% | 50.0% | 46.15% | 48.0% |
| **Decision Tree** | 52.56% | 66.33% | 52.0% | 52.56% | 50.0% | 66.67% | 38.46% | 57.14% |

AUROC = area under the receiver operating characteristic curve; AUPRC = area under the precision-recall curve; SVC = support vector classifier; NB = naive Bayes.
